# Autophagy regulates hyaluronan synthase 2 levels in vascular endothelial cells

**DOI:** 10.1101/413948

**Authors:** Carolyn G. Chen, Maria A. Gubbiotti, Xiaorui Han, Yanglei Yu, Robert J. Linhardt, Renato V. Iozzo

**Affiliations:** Department of Pathology, Anatomy and Cell Biology and the Cell Biology and Signaling Program, Kimmel Cancer Center, Sidney Kimmel Medical College at Thomas Jefferson University, Philadelphia, PA 19107, United States; Center for Biotechnology and Interdisciplinary Studies, Rensselaer Polytechnic Institute, Troy, NY 12180, United States

## Abstract

Hyaluronan is emerging as a key player regulating wound repair, inflammation, and angiogenesis. Of the three hyaluronan synthases (HAS1-3), HAS2 is the main driver of tumorigenicity by enhancing hyaluronan deposition. We discovered that HAS2 was degraded in vascular endothelial cells via autophagy, a catabolic process evoked by endorepellin and endostatin, two angiostatic and pro-autophagic effectors, and by Torin 1, a specific mTOR inhibitor. Protracted autophagy increased co-localization of HAS2 with three core components of the autophagosome, LC3, p62, and ATG9A, and binding of HAS2 to ATG9A. Importantly, autophagic induction led to an exclusive and marked suppression of secreted hyaluronan with no significant effects on either heparan or chondroitin sulfate levels. Thus, we have unveiled autophagy as a key catabolic mechanism regulating the production of hyaluronan in endothelial cells. Moreover, our study provides a biological link between autophagy and angiogenesis that could lead to potential targets for tumor neovascularization.

**Summary:** Hyaluronan is pro-angiogenic and pro-tumorigenic. We report a novel mechanism through which three autophagic inducers―endorepellin, endostatin, and Torin 1―regulate the levels of hyaluronan synthase-2. Protracted autophagy results in suppression of extracellular hyaluronan, thereby implicating autophagy in the regulation hyaluronan production.

## Introduction

The extracellular matrix (ECM) encompasses a variety of different signaling effectors to regulate an abundance of dynamic roles ranging from tissue homeostasis to development. Of note, extracellular proteoglycans masterfully synchronize a wide array of biological functions via outside-in cues (Iozzo, 2005), oscillating from maintaining tissue hydrodynamics and structural integrity to initiating cellular function and activating signal transduction via interaction with integrins and receptor tyrosine kinases (RTK) (Iozzo and Sanderson, 2011). In particular, the modular heparan sulfate proteoglycan (HSPG), perlecan, supports a host of functions, including cell adhesion, lipid metabolism, angiogenesis, endocytosis, thrombosis, and autophagy (Nugent et al., 2000; Iozzo and San Antonio, 2001; Ning et al., 2015; Poluzzi et al., 2014). Perlecan is present not only in vascular basement membranes and pericellular spaces (Gubbiotti et al., 2017), but also in human sera as a major circulating protein (Adkins et al., 2002). Structurally, it consists of a ∼500 kDa protein core spanning 5 domains, which undergoes partial proteolysis via matrix metalloproteinases to cleave its C-terminal domain V, endorepellin (Mongiat et al., 2003; Jung et al., 2013). Endorepellin consists of three laminin-like globular (LG) domains, allowing it to partake in partial antagonistic activity by dually signaling through the vascular endothelial growth factor receptor 2 (VEGFR2) and α2β1 integrin to induce protracted autophagy and disassemble the actin cytoskeleton, respectively. These contemporaneous actions of endorepellin ultimately lead to a robust anti-migratory and anti-angiogenic phenotype (Bix et al., 2004; Willis et al., 2013; Bix et al., 2007; Woodall et al., 2008). Importantly, endorepellin-mediated signaling via VEGFR2 also results in phosphorylation of adenosine monophosphate kinase (AMPK) at Thr^172^, leading to inhibition of mammalian target of rapamycin (mTOR) and induction of autophagy markers including Peg3, Beclin 1, LC3, and p62 (Torres et al., 2017; Buraschi et al., 2013; Liang et al., 1999; Yue et al., 2003; Goyal et al., 2014; Poluzzi et al., 2014). Taken together, these actions of endorepellin position it as a key soluble pro-autophagic and anti-angiogenic molecule with promising therapeutic effects against tumor vasculature (Mongiat et al., 2003; Bix et al., 2004; Bix and Iozzo, 2005; Bix et al., 2006; Douglass et al., 2015; Goyal et al., 2011; Goyal et al., 2012).

Hyaluronan (HA), another major component of the ECM, is a ubiquitous linear glycosaminoglycan composed of repeating disaccharide units of glucuronic acid and N-acetyl-glucosamine. HA is synthesized at the plasma membrane via multi-pass transmembrane enzymes of the HA synthase family (HAS1-3) (Weigel and DeAngelis, 2007). Each HAS exhibits unique glycosyltransferase activity that incorporates UDP-sugars to the non-reducing end of the growing glycosaminoglycan at the cytosolic leaflet of the membrane while simultaneously extruding the reducing end from the cell (Weigel and DeAngelis, 2007; Weigel, 2015). Although simple in composition, HA regulates a variety of cellular functions including wound repair, inflammation, cell migration, and angiogenesis (Oksala et al., 1995; Simpson, 2006; Simpson et al., 2015; Jackson, 2009) and recently emerged as a key player in regulating the tumorigenic and inflammatory milieu (Chanmee et al., 2016; Jackson, 2009). Intriguingly, its physiological effects are size-dependent: full-length HA (∼100-2000 kDa) is anti-inflammatory whereas its processed fragments (<36 kDa) are pro-inflammatory and pro-angiogenic (Jackson, 2009).

In a variety of diseases including cancer, HAS2 is the main driver of increased extracellular deposition of lower molecular weight HA, creating a microenvironment favoring angiogenesis and inflammation that perpetuates disease progression (Jackson, 2009; Chang et al., 2014). In breast cancer, aberrant overexpression of HAS2 has been shown to be involved in tumor growth and differentiation, axillary lymph node metastasis, and decreased patient survival (Auvinen et al., 2013; Auvinen et al., 2014; Bernert et al., 2011; Chanmee et al., 2016; Heldin et al., 2013; Karousou et al., 2017; Li et al., 2015b; Li et al., 2007). Furthermore, knockdown of HAS2 inhibits breast cancer growth and attenuates HA expression (Li et al., 2015b). Likewise, in other solid tumors of the ovary, colorectum, and prostate, HAS2 is an important regulator of tumorigenicity and metastasis through excessive production of HA (Chanmee et al., 2016). Notably, phosphorylation of HAS2 by AMPK and *O*-GlcNAcylation modulate its enzymatic activity and stability, respectively (Vigetti et al., 2014; Vigetti et al., 2011a).

As endorepellin evokes autophagy via AMPK (Poluzzi et al., 2014; Poluzzi et al., 2016), an enzyme critical for autophagic progression (Garcia and Shaw, 2017; Herzig and Shaw, 2018), and as AMPK regulates HAS2 activity (Vigetti et al., 2011b), we hypothesized that HAS2 levels could also be regulated via autophagy. Here, we leveraged three independent activators of autophagy to investigate the modulation of protracted autophagy on HAS2: endorepellin, the proteolytic fragment of perlecan (Goyal et al., 2016; Douglass et al., 2015), Torin 1, an ATP-competitive second generation mTOR inhibitor (Thoreen et al., 2009), and endostatin, the C-terminal fragment of collagen XVIII (Nguyen et al., 2009; Poluzzi et al., 2016). We showed that all three activators evoked downregulation and autophagic flux of HAS2 protein, with the former two resulting in a selective and marked decrease in extracellular HA. Additionally, we demonstrated that HAS2 bound ATG9A, the only multi-pass transmembrane protein in the core autophagic complex, and that this interaction was enhanced with autophagic activation. Together, these findings elucidate a universal mechanism of HAS2 regulation activated by ECM-mediated signaling to suppress synthesis of secreted HA, which may ultimately be used to craft novel chemotherapeutic strategies that target the tumor stroma.

## Results and discussion

### Endorepellin downregulates HAS2 in endothelial cells

To investigate the intracellular catabolism of HAS2, we utilized endorepellin, the potent pro-autophagic C-terminus of perlecan. Remarkably, we found that soluble endorepellin downregulated HAS2 levels in a dose-dependent fashion in vascular endothelial cells (IC_50_∼270 nM, Fig. 1, A and B). Concurrently, endorepellin downregulated VEGFR2 activity by reducing phosphorylation at Tyr^1175^ with a concomitant activation of AMPKα by phosphorylation at Thr^172^ (Fig. 1, A-D). Furthermore, endorepellin downregulated HAS2 expression in a time-dependent fashion with a calculated T_1/2_ of 7.5 h (Fig. 1, E and F) while inhibiting P-VEGFR2 activation at Tyr^1175^ and increasing P-AMPKα activation at Thr^172^ (Fig. 1, G and H). Mechanistically, transient knockdown of VEGFR2 via siRNA ablated the downregulation of HAS2 by endorepellin (Fig. 1, I-K), indicating that endorepellin signals through VEGFR2 to downregulate HAS2. Notably, this modulation of HAS2 occurred post-transcriptionally, as mRNA profiles of *HAS2* determined via quantitative real-time PCR (qPCR) remained unchanged in HUVEC treated with endorepellin for 6 and 24 h vis-à-vis vehicle (Fig. S1 A). Moreover, *HAS3* showed no significant changes in the presence of endorepellin (Fig. S1 B), whereas *HAS1* levels were barely detectable (Fig. S1 C). Thus, the pro-autophagic perlecan fragment endorepellin dynamically downregulates HAS2 in endothelial cells at the protein level.

**Figure 1.**
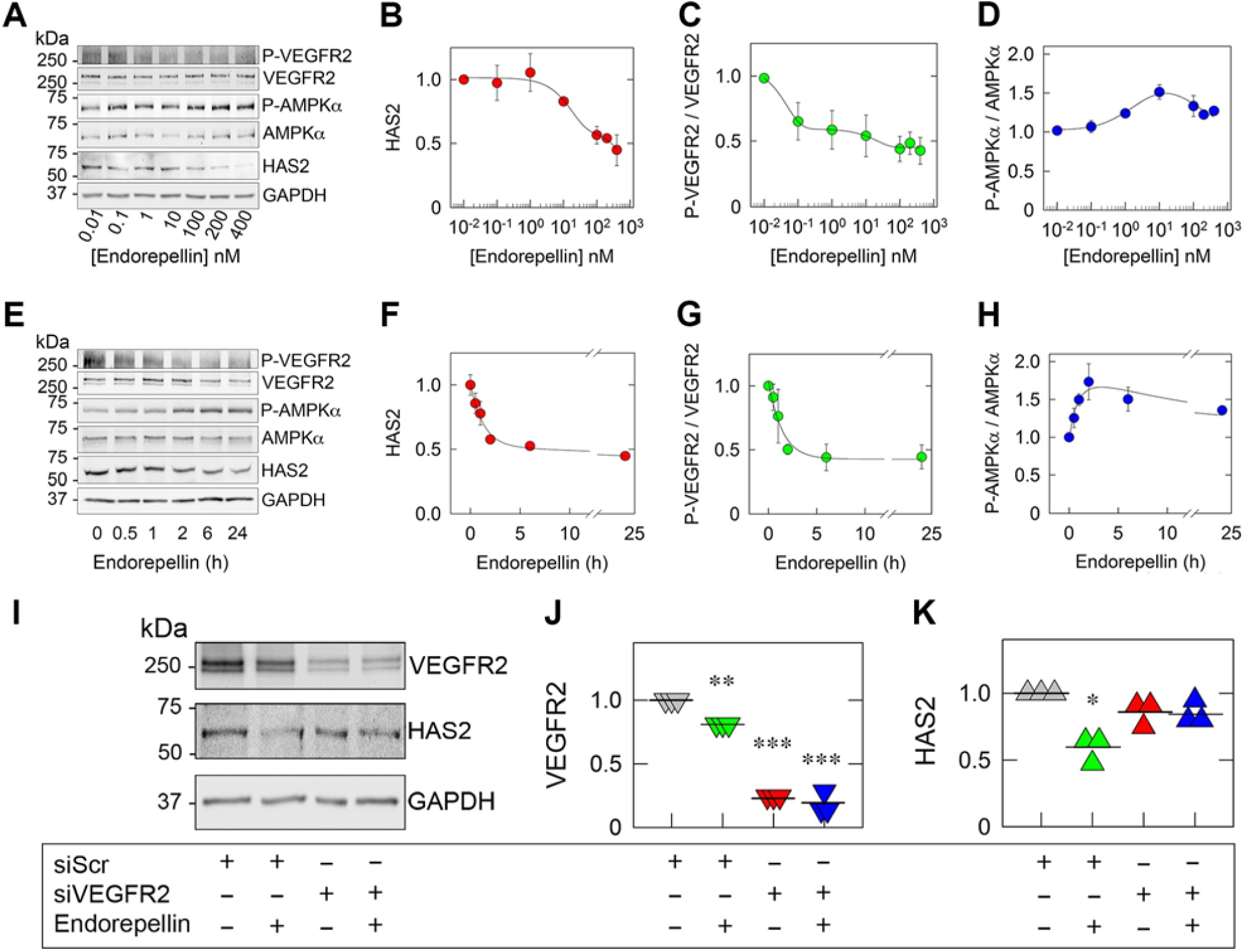
Endorepellin and endostatin downregulate levels of HAS2 protein. **(A)** Representative western blot (WB) showing downregulation of HAS2, downregulation of P-VEGFR2^Y1175^, and induction of P-AMPKα^T172^ in HUVEC whole-cell lysates treated with increasing concentrations of endorepellin for 6 h. **(B-D)** Quantification of HAS2, P-AMPKα^T172^ and P-VEGFR2^Y1175^, respectively, from **(A)**. **(E)** Representative WB of HUVEC whole-cell lysates treated with endorepellin (200 nM) at increasing time points. **(F-H)** Quantification of HAS2, P-VEGFR2^Y1175^ and P-AMPKα^T172^, respectively, from **(E)**. Means ± SE, n = 3 biological replicates. Data are normalized to GAPDH. **(I)** Representative immunoblots of HUVEC pretreated with scrambled siRNA (siScr) or siRNA targeting VEGFR2 (siVEGFR2) following treatment with endorepellin (200 nM) for 6 h. **(J-K)** Quantification of VEGFR2 and HAS2 from **(I)** (n=3). Data are normalized to GAPDH and presented as the mean (horizontal bar). Statistical analyses were calculated via Student’s t test. *, P < 0.05; **, P < 0.01; ***, P < 0.01.

### The selective mTOR inhibitor Torin 1 downregulates HAS2 in endothelial cells

To validate that endorepellin-mediated degradation of HAS2 is autophagic in nature, we evaluated HAS2 levels in HUVEC following treatment with Torin 1, a competitive mTOR inhibitor. Torin 1 canonically induces autophagy by directly binding to the active site of mTOR, making it a potent and specific autophagic inducer (Thoreen et al., 2009). Similar to endorepellin, Torin 1 evoked a dose-dependent suppression of HAS2 protein (IC_50_∼188 nM, Fig. 2, A and B). We verified induction of autophagy by noting a decrease of autophagic marker, p62, (Fig. 2 C) and concurrent induction of lipidated LC3-II, a well-characterized component of the autophagosomal membrane (Fig. 2 D) (Mizushima et al., 2010; Komatsu and Ichimura, 2010). We then treated HUVEC with Torin 1 over 24 h and observed an analogous effect where Torin 1 (20 nM) downregulated HAS2 (T_1/2_∼5.2 h, Fig. 2, E and F). Likewise, p62 levels were diminished and LC3-II levels were induced with increasing treatment times of Torin 1. Additionally, endostatin, the pro-autophagic C-terminal fragment of collagen XVIII shown to inhibit mTOR expression (Wang et al., 2015), also downregulated HAS2 in a dose-dependent manner while upregulating LC3-II (Fig. 2, I-K), further implicating autophagy as a general pathway of regulation for HAS2. Thus, protracted autophagy via mTOR inhibition effectively downregulates HAS2 in endothelial cells.

**Figure 2.**
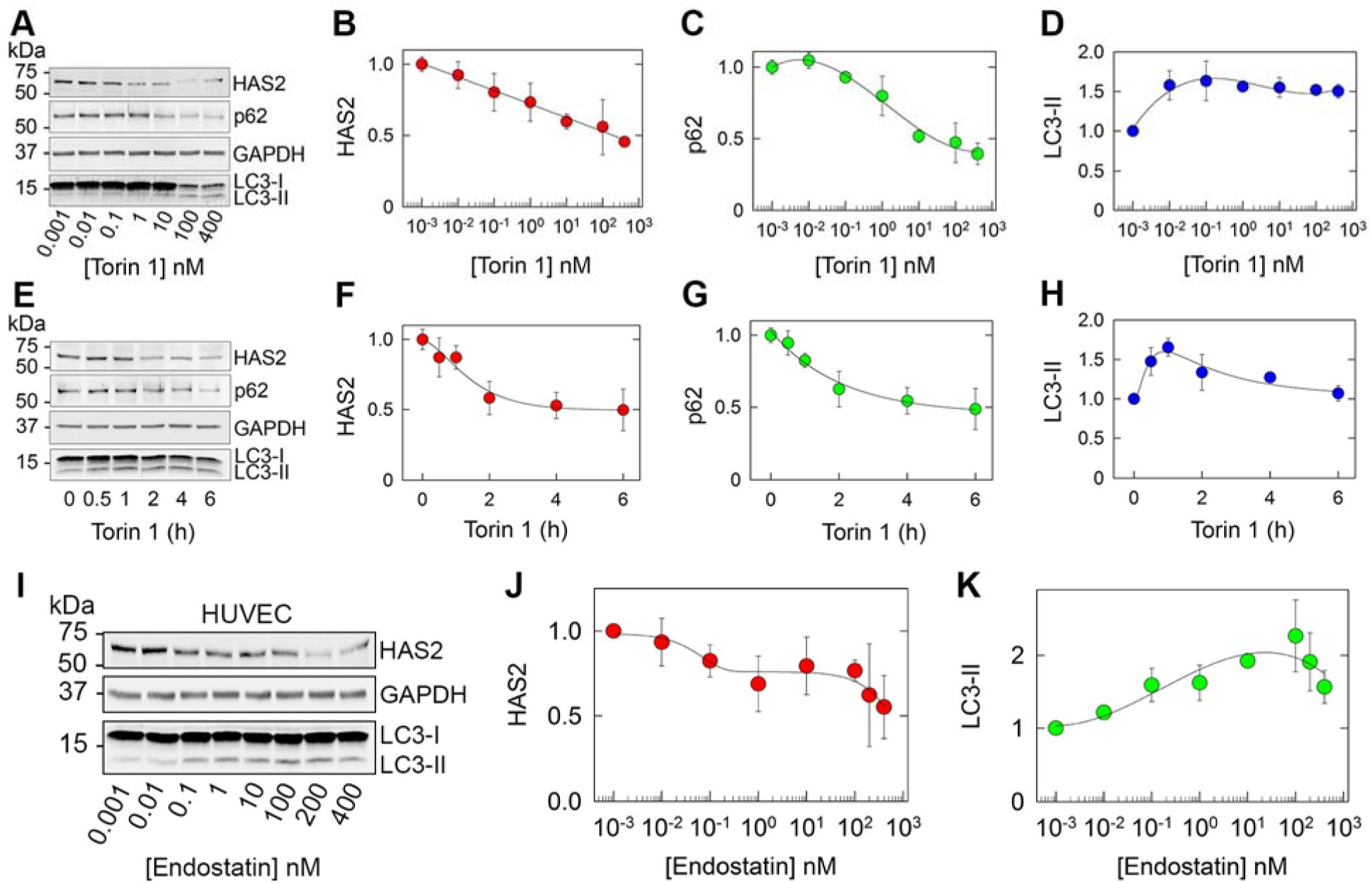
mTOR inhibition via Torin 1 and endostatin downregulate levels of HAS2 protein. **(A)** Representative WB showing downregulation of HAS2, downregulation of p62, and induction of LC3-II in HUVEC whole-cell lysates treated with increasing concentrations of Torin 1 for 6 h. GAPDH served as loading control. **(B-D)** Quantification of HAS2, p62 and LC3-II, respectively, from **(A)**. Means ± SE presented as the mean (horizontal bar); n = 3 biological replicates. **(E)** Representative WB of HUVEC whole-cell lysates treated with Torin 1 (20 nM) at increasing time points. **(F-H)** Quantification of HAS2, p62 and LC3-II, respectively, from **(E)**. Means ± SE, n = 3 biological replicates. Data are normalized to GAPDH. **(I)** Representative WB showing decreased HAS2 in HUVEC whole-cell lysates treated with increasing concentrations of endostatin for 6 h. GAPDH served as the loading control. Quantification of HAS2 **(J)** and LC3-II **(K)** normalized to GAPDH from **(L)**. Data are presented as means ± SE (n = 3).

### HAS2 co-localizes with LC3, p62, and ATG9A in autophagosomal structures

Having established a link between endorepellin/mTOR and autophagy, we next tested whether HAS2 would co-distribute with members of the autophagosomal pathway. Under basal conditions, LC3 is localized in the cytosol; however, during autophagy, LC3 becomes lipidated to form LC3-II and localizes to the autophagosomal membrane (Mizushima et al., 2010). Similarly, p62, also referred to as sequestosome 1 (SQSTM1), is correspondingly degraded by autophagy and binds simultaneously to both ubiquitinated proteins and LC3 and GABARAP family proteins (Komatsu and Ichimura, 2010). To this end, we utilized Bafilomycin A1, a pharmacological inhibitor of autophagosome-lysosome fusion that blocks activity of vacuolar (V-type) H^+^- ATPases (Levine and Kroemer, 2008). Confocal imaging showed that HAS2 co-localized with both LC3 and p62 under basal conditions (Fig. 3, A and B), and this co-localization was markedly enhanced when autophagic flux was blocked by Bafilomycin A1 (Fig. 3, A and B, insets). Next, we investigated the interaction between HAS2 and ATG9A, both of which are multi-pass transmembrane proteins. Notably, ATG9A plays a key role in nucleating the pre-autophagosomal structure (PAS), cycling between the trans-Golgi network and endosomes and contributing plasma membrane to the developing autophagosome (Mattera et al., 2017; Zhuang et al., 2017). We discovered that co-localization of HAS2 and ATG9A upon Bafilomycin A1 treatment was increased vis-à-vis vehicle (Fig. 3 C), implicating a potential role for HAS2 to subsidize membrane components to the growing autophagosome. To verify this co-localization, we utilized line scanning and quantified the relative pixel intensity of differentially labeled fluorophores along a single axis in autophagosomal structures (Thurston et al., 2012; Sai et al., 2008). All three autophagic markers displayed a high degree of overlap, indicating extensive co-localization with HAS2 (Fig. 3, A-C). Thus, we conclude that HAS2 is an autophagic substrate as it co-localizes with all three autophagic markers LC3, p62, and ATG9A in cytoplasmic puncta.

**Figure 3.**
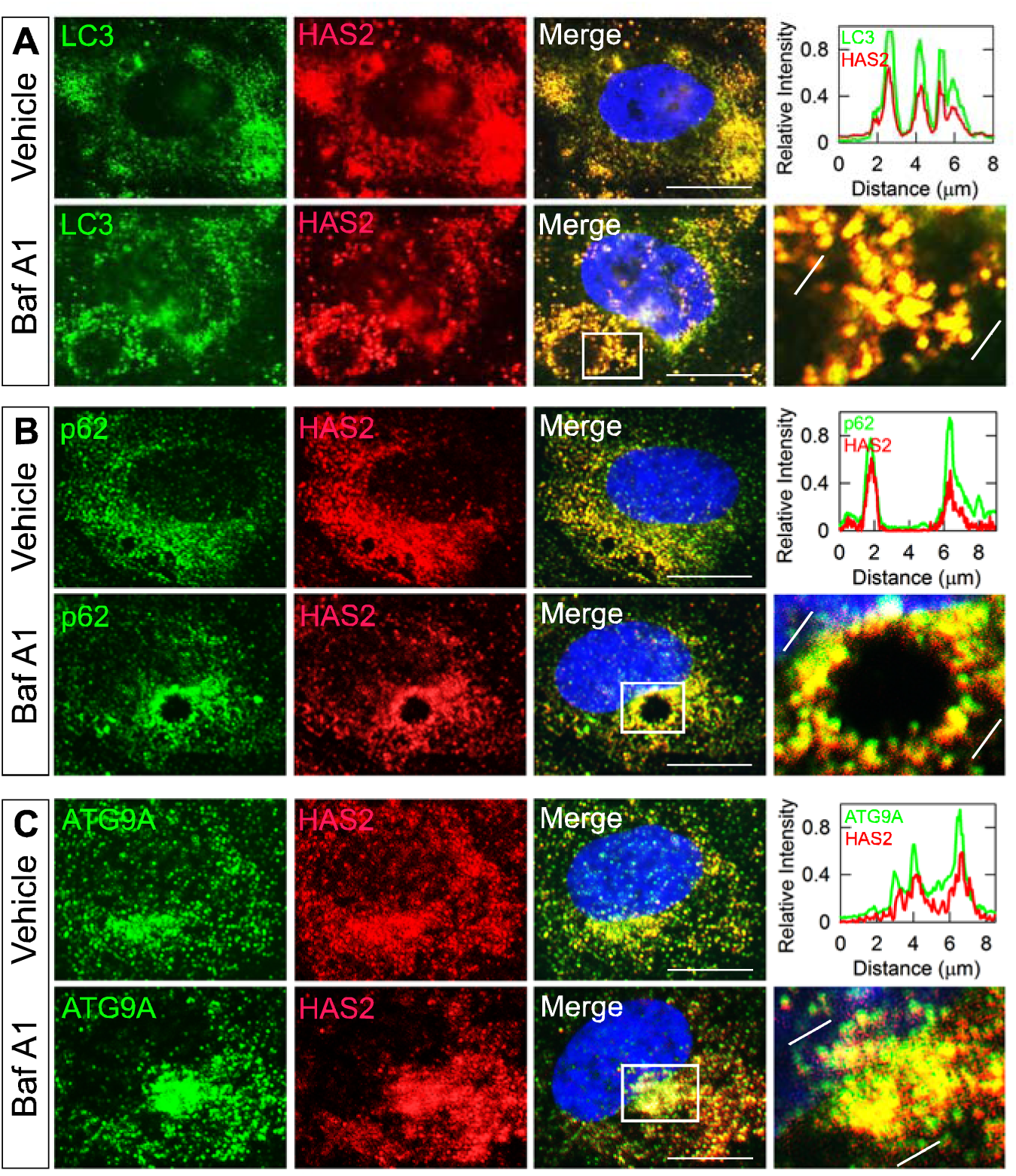
Bafilomycin A1 induces co-localization of HAS2 with autophagosomal proteins LC3, p62 and ATG9A. Representative immunofluorescent confocal images of HUVEC following treatment with vehicle (DMSO) or Bafilomycin A1 (500 nM; abbreviated as Baf A1) for 6 h. The cells were dually labeled for HAS2 (red) and either LC3 **(A)**, p62 **(B)**, or ATG9A (green) **(C)**. In the *Right* column, the bottom and top portions of each panel are enlarged insets of the white boxed areas and their corresponding line-scanning profiles, respectively. Line-scanning profiles taken between the white lines measure the fluorescence intensity for HAS2 (red lines) and LC3, p62, or ATG9A (green lines) (blue, DAPI). Scale bar, 10 μm.

### Endorepellin and Torin 1 evoke autophagic flux of HAS2 and co-localization with LC3 and ATG9A

To verify that HAS2 expression was modulated via autophagy, we performed autophagic flux assays with Bafilomycin A1 to measure the build-up of autophagic substrates following provocation of this catabolic process. Western blot analyses of HUVEC lysates showed significantly increased levels of HAS2 with Bafilomycin A1 vis-à-vis vehicle (Fig. 4, A and B), with a concurrent accumulation of p62 and LC3-II (Fig. 4, C and D). Notably, compared to Bafilomycin A1 alone, addition of either endorepellin or Torin 1 caused a significant build-up of HAS2, p62 and LC3-II (Fig. 4, A-D), verifying effective autophagic induction.

**Figure 4.**
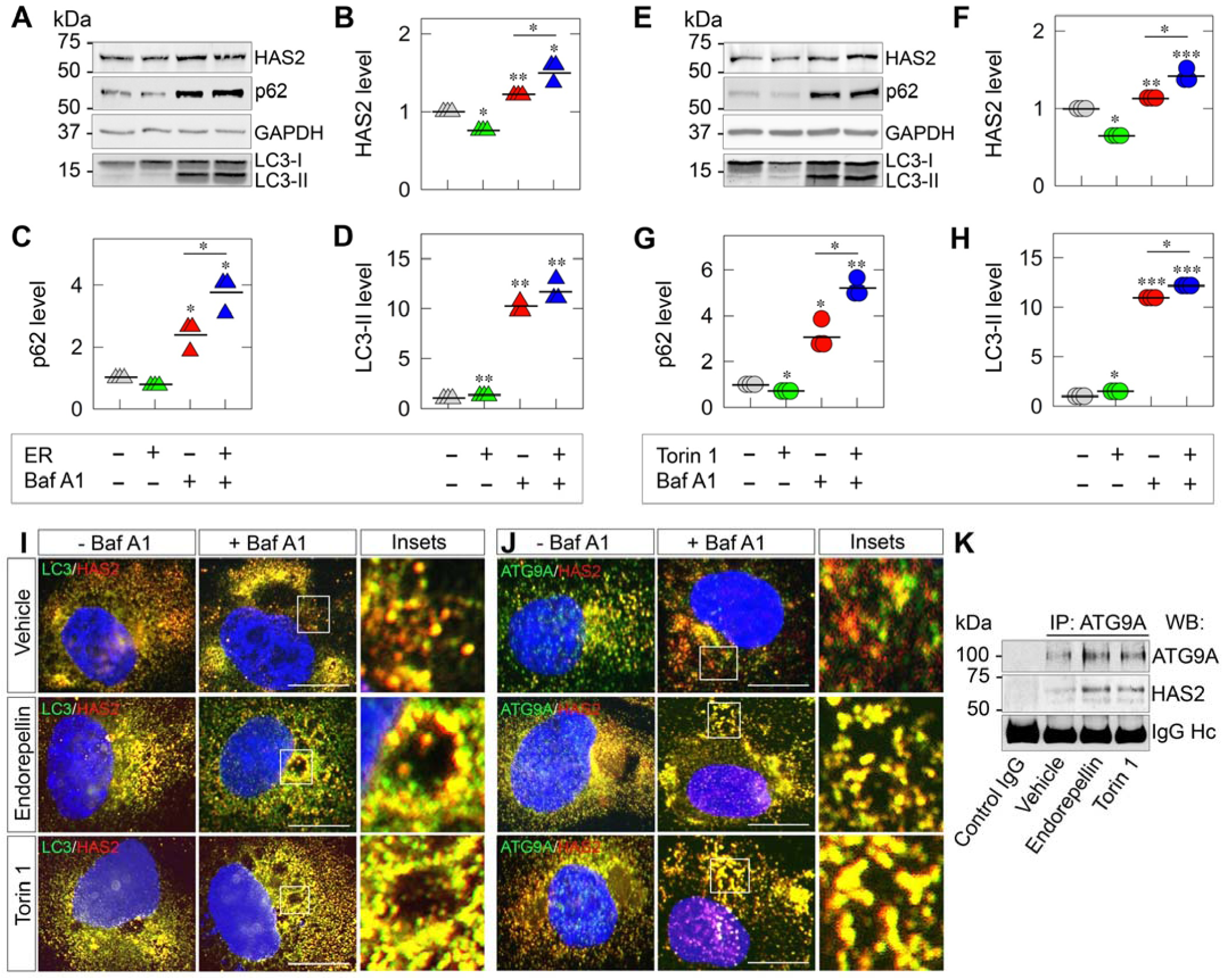
Endorepellin and Torin 1 separately induce autophagic flux and co-localization of HAS2 into LC3- and ATG9A-positive autophagosomes. **(A)** Representative WB of HAS2, p62, LC3, and GAPDH following treatment with vehicle or endorepellin (200 nM, 6 h; abbreviated as ER) ±Bafilomycin A1 (500 nM; abbreviated as Baf A1). **(B-D)** Quantification of HAS2, p62, and LC3-II, respectively, from **(A)** (n=3). **(E)** Corresponding confocal images of HUVEC following treatment in **(A)** and dually labeled for HAS2 (red) and either LC3, p62, or ATG9A (green). **(F)** Representative WB of HAS2, p62, LC3, and GAPDH following treatment with vehicle or Torin 1 (20 nM, 6 h) ±Bafilomycin A1 (500 nM). **(G-I)** Quantification of HAS2, p62, and LC3-II, respectively, from **(F)** (n=3). Data are normalized to GAPDH and presented as the mean (horizontal bar). Statistical analyses were calculated via Student’s t test. *, P < 0.05; **, P < 0.01; ***, P < 0.01. **(J)** Corresponding confocal images of HUVEC following treatment in **(F)** and dually labeled for HAS2 (red) and either LC3 or ATG9A (green). Enlarged insets depict co-localized (yellow) signal from cells treated with Bafilomycin A1 and endorepellin or Torin 1 (Right). Scale bar, 10 μm. **(K)** Representative co-immunoprecipitation (co-IP) of HUVEC stimulated with endorepellin (200 nM) or Torin 1 (20 nM) for 3 h. The cells were lysed, immunoprecipitated (IP) with an anti-ATG9A antibody and subjected to WB with anti-ATG9A and anti-HAS2 antibodies as indicated. IgG heavy chain (Hc) is presented as a loading control.

Next, we corroborated these biochemical changes with confocal imaging which showed enhanced co-localization of HAS2 with LC3 in cells treated with endorepellin or Torin 1 ±Bafilomycin A1 (Fig. 4 I). Compared to vehicle, endorepellin or Torin 1 treatment evoked increased co-localization of HAS2 and LC3 into autophagosomal vacuoles, an effect that was strongly enhanced in the presence of Bafilomycin A1, with insets depicting punctate co-localization of HAS2 and LC3. A similar trend of co-localization of HAS2 and p62 was observed in HUVEC (Fig. S2 A). Importantly, confocal microscopy demonstrated enhanced co-localization of HAS2 with ATG9A in autophagosomal structures evoked by either endorepellin or Torin 1 vis-à-vis vehicle (Fig. 4 J). As a quantitative measure, we calculated the interaction of HAS2 with ATG9A, LC3, and p62 using the Pearson’s coefficient of co-localization for overlapped pixels (Misaki et al., 2010; Dunn et al., 2011; Zinchuk et al., 2011). Upon treatment with endorepellin or Torin 1 for 6 h, the degree of co-localization between HAS2 and ATG9A, LC3, or p62 was significantly increased vis-à-vis vehicle (Fig. S2, B-D). Finally, we were able to co-immunoprecipitate complexes of ATG9A and HAS2, detectable under basal conditions and markedly enhanced by endorepellin or Torin 1 (Fig. 4 K). In contrast, we were unable to co-immunoprecipitate HAS2 with either LC3 or p62 (Fig. S3). Collectively, our results demonstrate that HAS2 is a substrate of autophagic flux and open the possibility of a physical interaction between two crucial multi-pass transmembrane proteins in endothelial cells.

### Endorepellin and Torin 1 selectively suppress extracellular HA

Having established that induction of autophagy by various agents leads to suppression of HAS2, and given the fact that this enzyme is a major producer of HA (Iozzo and Schaefer, 2015), we measured the glycosaminoglycan composition of media conditioned by endothelial cells. We performed quantitative disaccharide analyses with liquid chromatography-tandem mass spectrometry (LC-MS/MS) using defined unsaturated disaccharide standards (Table S1) and Multiple Reaction Monitoring (MRM) (Table S2) (Li et al., 2015a). Relative to the vehicle, HA secreted from cells treated with endorepellin and Torin 1 had a notable 3- and 4.33-fold decrease, respectively (p<0.01) (Fig. 5 A). In contrast, protracted autophagy by endorepellin or Torin 1 had no significant effects on extracellular HS, CS, or total GAG levels (Fig. 5, B and C). Thus, endorepellin and Torin 1 specifically target this important enzyme for intracellular degradation without appreciably affecting the biosynthesis of other glycosaminoglycans.

**Figure 5.**
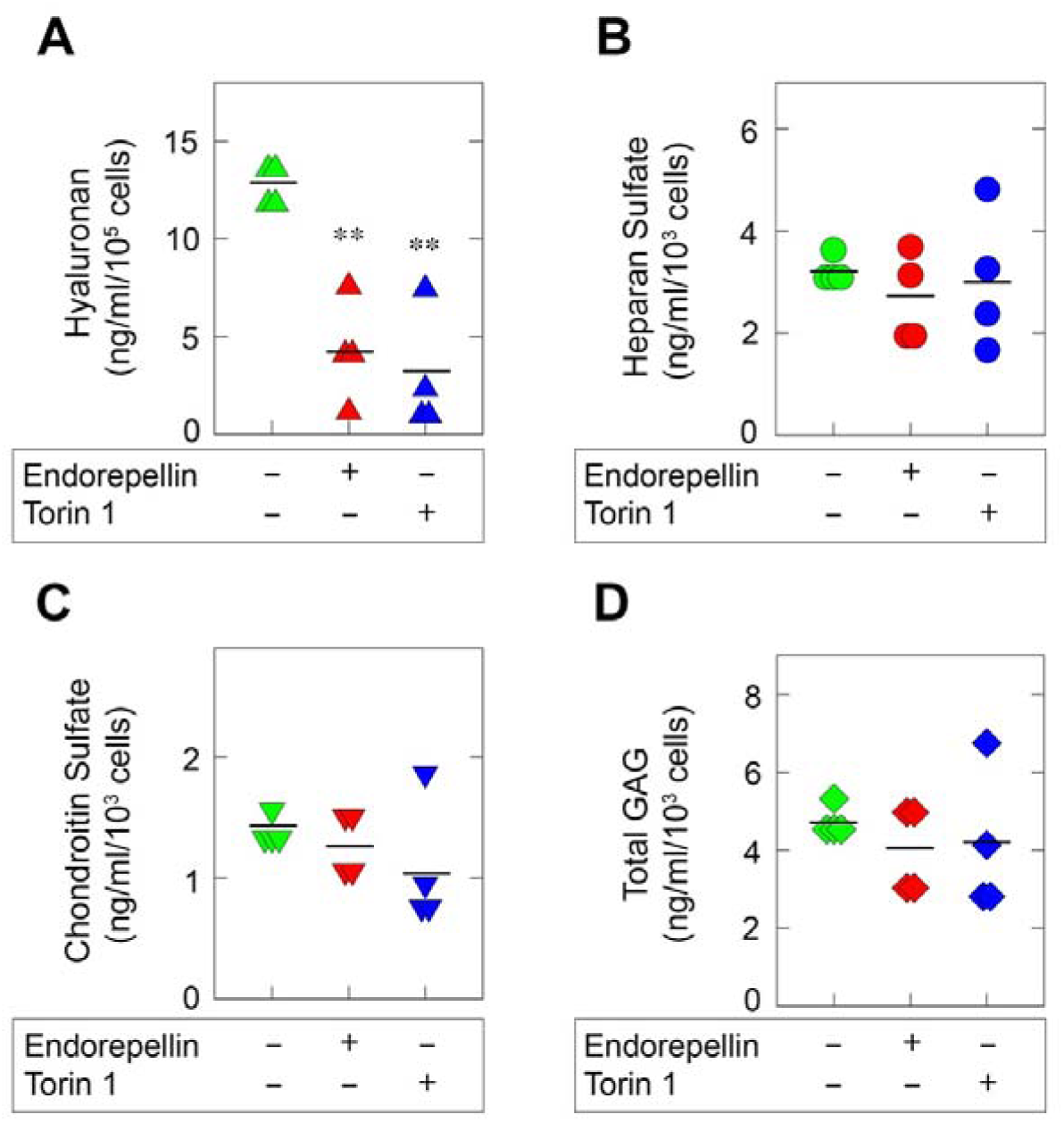
Endorepellin and Torin 1 separately suppress secreted HA with no significant effect on heparan sulfate (HS), chondroitin sulfate (CS), and total glycosaminoglycan (GAG) levels. Quantification by liquid chromatography-mass spectrometry (LC-MS) of total secreted HA **(A)**, HS **(B)**, CS **(C)**, and total GAG **(D)** in HUVEC media conditioned for 24 h by vehicle, endorepellin (200 nM) or Torin 1 (20 nM). Data are normalized to GAPDH and presented as the mean (horizontal bar); n=4. Statistical analyses were calculated via Student’s t test. **, P < 0.01

Unlocking the secrets of the tumor stroma has expanded our understanding of the intricate processes of cancer initiation and progression. Particularly, dynamic remodeling of the ECM is becoming increasingly acknowledged as an oncologic and angiogenic force, as tumor cells and their vascularized stroma exert paracrine and angiocrine modes of regulation to stimulate angiogenesis, cell growth, and metastasis (Bissell and Radisky, 2001; Bissell and Hines, 2011; Quail and Joyce, 2013; Butler et al., 2010). Over the years, our lab has provided solid evidence that the ECM component, endorepellin, evokes an anti-angiogenic response *ex vivo* and *in vivo* (Goyal et al., 2016; Bix and Iozzo, 2008) by targeting the tumor vasculature, thereby inhibiting tumor angiogenesis and growth and increasing tumor hypoxia in squamous and Lewis lung carcinoma (Bix et al., 2006). Since the cognate receptor for endorepellin, VEGFR2, is not expressed on the surface of these tumor cells, endorepellin’s anti-cancer activity is solely due to its ability to influence signaling in the tumorigenic milieu, thereby underscoring the importance and need for more in-depth study of the modulation of tumorigenesis via stromal signaling evoked by ECM effectors. Moreover, the most recent focus of endorepellin’s biologic activity in endothelial cells involves its ability to induce protracted autophagy, a physiologic housekeeping process that, in moderation, serves to eliminate damaged organelles, clear protein aggregates, and maintain cellular homeostasis (Yang and Klionsky, 2010; Moresi et al., 2008). However, a significant portion of the deranged signaling seen in malignant processes can be attributed to aberrations of autophagy both in the tumor parenchyma and supporting cells (Galluzzi et al., 2017; Levine and Kroemer, 2008). Importantly, suppression of a pro-oncogenic EGFR via RTK inhibitor therapy induces autophagy in non-small-cell lung carcinoma (Wei et al., 2013), whereas enhanced autophagy blocks HER2-mediated breast tumorigenesis (Vega-Rubin-de-Celis et al., 2018). However, despite these significant scientific contributions, no pharmacologic interventions specifically targeting autophagy are currently clinically available for human use to confront cancer (Galluzzi et al., 2017; Rubinsztein et al., 2012).

HAS1-3 isoforms comprise the family of HA synthases that polymerize HA in the intracellular face of the plasma membrane. Each of these enzymes polymerize varying chain lengths of HA, with both HAS1 and HAS2 producing high molecular weight HA (∼2000 kDa) and HAS3 producing lower molecular weight HA (100-1000 kDa). Under physiological conditions, most of the extracellular HA is of relatively high molecular weight, whereas HA polymers are significantly fragmented into oligosaccharides during tumorigenesis and inflammation, evoking a pro-angiogenic and pro-inflammatory phenotype (Karousou et al., 2017). In the present work, we identify a novel mechanism to regulate pro-angiogenic HA via targeting HAS2 in vascular endothelial cells. A previous study showed modification of HAS2 with polyubiquitin chains linked at the Lys-48 residue, the linkage selectively targeted for degradation to proteasomes (Karousou et al., 2010). However, blocking proteasome-mediated degradation via MG132 only slightly increased the pool of polyubiquitinated HAS2 and did not significantly alter total HAS2 nor its capacity to synthesize HA (Karousou et al., 2010). We now provide robust evidence that HAS2 protein levels are regulated by canonical autophagy. Mechanistically, all three autophagic inducers endorepellin, endostatin, and Torin 1, independently activate the canonical autophagic pathway by converging on mTOR inhibition (Goyal et al., 2012; Wang et al., 2015). Indeed, we previously discovered that endorepellin mobilizes mTOR into LC3-positive autophagosomes in endothelial cells (Goyal et al., 2016). In support of our findings, an independent study has found that mTOR activation in fibroblasts increases levels of HAS2 and secreted HA, and that this effect could be partially reversed by rapamycin, an indirect inhibitor of mTOR (Qin et al., 2012). However, this study did not explore autophagy as a mechanism of HAS2 regulation downstream of mTOR inhibition. Thus, our study provides evidence to support an mTOR-dependent autophagic regulation of this enzyme. Furthermore, the discovery of autophagic clearance of HAS2 and subsequent downregulation of HA via mTOR inhibition is generalizable in a broad context given the various approaches of mTOR modulation and autophagic induction.

The interaction between two multi-pass transmembrane proteins, one involved in in HA synthesis (HAS2) and another (ATG9A) contributing to the pre-autophagosomal structure, is interesting and suggests the possibility that HAS2 and ATG9A could co-distribute perhaps within lipid rafts or specialized endocytic vesicles. Notably, ATG9A is crucial for providing contact sites between the endoplasmic reticulum and the phagophore (Gomez-Sanchez et al., 2018), thereby promoting formation of the autophagosomal membrane. Other studies suggest the trans-Golgi network and plasma membrane can contribute to the autophagosomal membrane (Mattera et al., 2017; Pavel and Rubinsztein, 2017; Levine, 2016). As an integral membrane protein, HAS2 could play a mechanistic role in forming the pre-autophagosomal structure, perhaps by shuttling lipid components to the growing autophagosome via complexing with ATG9A upon autophagic activation. This may be a function of HAS2 independent of its enzymatic activity, a notion requiring further investigation.

In this study, we show that autophagic clearance of HAS2 results in a marked depletion of secreted HA with no changes in other GAGs, specifically HS or CS. Since HA shares an overlapping pool of monosaccharide substrates with HS and CS, the selective reduction in HA is likely not due to depleted cytosolic stores of glucuronic acid or N-acetyl-glucosamine, the building blocks of HA (Volpi and Linhardt, 2010), but rather the targeted depletion of HAS2. Furthermore, recent studies have implicated the physiologically active forms of HAS as homo- or heterodimers (Vigetti et al., 2014; Karousou et al., 2010), with HAS2 playing a critical role in the formation of enzymatically active HAS2 or HAS3 homodimers and HAS2/HAS3 heterodimers. Thus, downregulating HAS2 may affect overall HA synthesis by disrupting the cooperative dimerization of these synthases. These data present an autocrine and paracrine mode of regulation in vascular endothelial cells where an intracellular degradative pathway alters a distinct component of the secretome that, in turn, suppresses pro-angiogenic signals to the cell, leading ultimately to an anti-angiogenic phenotype. Thus, we propose that selectively targeting HA by HAS2-modulation via the induction of autophagy is a promising avenue of investigation for chemotherapeutics as HA is a major proponent of cancer cell angiogenesis, proliferation, and metastasis (Nagy et al., 2015).

Taken together, these results shift the current archetype of autophagy from a mere quality control process to one that precisely regulates the expression of key ECM constituents affecting cancer progression and invasion (Choi et al., 2013; Mizushima and Levine, 2010). We believe that these findings will forge new paths to build an arsenal of powerful and effective cancer treatments that augment autophagy and regulate its downstream targets, especially in the tumor microenvironment, resulting in medical advances to better serve cancer patients.

## Materials and Methods

### Cells and materials

Human umbilical vein endothelial cells (HUVEC) were grown and maintained in basal media supplemented with VascuLife EnGS LifeFactors Kit (Lifeline Cell Technology, Oceanside, CA). Cells were plated on 0.2% gelatin-coated cell culture plates (Thermo Fisher Scientific, Waltham, MA) and utilized within the first five passages. Antibodies were purchased from the following sources and utilized at the designated dilutions: GAPDH (14C10, 1:10,000 dilution; Cell Signaling, Danvers, MA), VEGFR2 (55B11, 1:1,000 dilution; Cell Signaling), P-VEGFR2 (19A10, 1:750 dilution; Cell Signaling), AMPKα (2532, 1:1,000 dilution; Cell Signaling), P-AMPKα (40H9, 1:1,000 dilution; Cell Signaling), ATG9A [ab108338, used for western blot (WB) at 1:1,000 dilution, immunofluorescence (IF) at 1:200, and immunoprecipitation (IP) at 4 μg; Abcam, Cambridge, MA], p62/SQSTM1 (ab91526, used for WB at 1:1,000 dilution and IF at 1:200; Abcam), p62/SQSTM1 (P0068, IP at 4 μg; Sigma-Aldrich, St. Louis, MO), LC3B (L7543, used for WB at 1:1,000 dilution and IF at 1:200; Sigma), HAS2 (sc-514737, used for WB at 1:250; Santa Cruz Biotechnology, Dallas, TX), HAS2 (sc-34068, used for IF at 1:50; Santa Cruz Biotechnology), and normal rabbit IgG (2729, used for IP at 4 μg; Cell Signaling). Secondary antibodies for IF were purchased from the following sources and utilized at the designated dilutions: Alexa Fluor 594 donkey anti-goat H+L IgG (A11058, 1:400; Thermo Fisher) and Alexa Fluor 488 goat anti-rabbit H+L IgG (A11008, 1:400; Thermo Fisher). The following siRNA was purchased from Santa Cruz Biotechnology and used at the designated amounts: Flk1-1 siRNA (sc-9318, 100 pM) and scramble control siRNA-A (sc-37007, 100 pM). Torin 1 and Bafilomycin A1 were purchased from Sigma. Protein A Sepharose magnetic beads were purchased from GE Healthcare. Recombinant human endorepellin was produced and purified as described previously (Mongiat et al., 2003). Murine Fc-Endostatin (1 mg/ml) was obtained from the NCI-BRB Preclinical Repository (Frederick, MD).

### Immunoblotting and immunoprecipitation

Treated HUVEC were rinsed once in ice-cold phosphate-buffered saline (PBS) and lysed in radioimmune precipitation assay (RIPA) buffer [50 mM Tris (pH 7.4), 150 mM NaCl, 1% Triton X-100, 0.1% sodium deoxycholate, 0.1% SDS, 1 mM EDTA/EGTA/sodium orthovanadate, 10 mMβ-glycerophosphate, and protease inhibitors (1 mM phenylmethylsulfonyl fluoride and 10 μg/ml leupeptin/tosylphenylalanyl chloromethyl ketone/aprotinin each)] for 15 min rocking on ice. Lysates were boiled for 3 min and resolved on SDS-PAGE. Proteins were then transferred to nitrocellulose membranes (Bio-Rad, Hercules, CA), probed with primary and secondary antibodies, and developed with SuperSignal West Pico Chemiluminescence substrate (Thermo Fisher Scientific). Signal detection was performed via an ImageQuant LAS-4000 (GE Healthcare) visualization platform.

For immunoprecipitation, protein A Sepharose magenetic beads (GE Healthcare) were incubated with antibodies for 4 h at room temperature (RT), and precleared cell lysates were added to the bead-antibody conjugate for 18 h at 4 °C. After extensive washing, beads were vortexed at high speed in reducing buffer for 30 min at RT (ATG9A IP) or boiled for 5 min (p62 IP), then separated using magnets. Visualizing ATG9A using the anti-ATG9A antibody was most clear in non-boiled lysates, as recommended by the manufacturer (Abcam). Remaining proteins in reducing buffer were analyzed via WB.

### qPCR analysis

Transcriptional expression analysis by real-time quantitative PCR (qPCR) was carried out on confluent 12-well plates with ∼2 × 10^5^ HUVEC treated with endorepellin (200 nM) or Torin 1 (20 nM) for 6 and 24 h in full serum Basal Endothelial Medium (Lifeline Cell Technology). Following incubation, cells were lysed in 500 μl TRIzol reagent (Invitrogen) to extract total RNA. Then, 1 μg of total RNA was annealed with oligo (dT_18-20_) primers, and SuperScript Reverse Transcriptase III (Thermo Fisher Scientific) was utilized to synthesize cDNA. Amplicons representing target genes and the endogenous housekeeping gene, β-2-microglobulin (*B2M*), were amplified in quadruple independent reactions using the Brilliant SYBR Green Master Mix II (Agilent Technologies, Santa Clara, CA). All samples were run on the Roche LightCycler 480-II Real Time PCR platform (Roche Applied Sciences, Indianapolis, IN), and cycle number (Ct) was recorded for each reaction. Fold changes were normalized to *B2M* or *HPRT1* and calculated using the ΔΔCt method (2^-^ΔΔCT). Data were derived from three independent biological replicates, each carried out in quadruplicate for every gene of interest. The primers used in qPCR analyses were as follows: *HAS1* forward, 5*’*- CTTGTCAGAGCTACTTCCACTG-3*’*; *HAS1* reverse, 5*’*-CGGTCATCCCCAAAAGTACAG-3*’*; *HAS2* forward, 5*’*- ATGGTTGGAGGTGTTGGG-3*’*; *HAS2* reverse, 5*’*-AGGTCCACTAATGCACTGAAC-3*’*; *HAS3* forward, 5*’*-atcatgcagaagtggggagg-3*’*; *HAS3* reverse, 5*’*-tccaggactcgaagcatctc-3*’*; *B2M* forward, 5*’*- TCCATCCGACATTGAAGTTG-3*’*; *B2M* reverse, 5*’*-ACACGGCAGGCATACTCAT-3*’*; *HPRT1* forward, 5*’*- GCTATAAATTCTTTGCTGACCTGCTG-3*’*; *HPRT1* reverse, 5*’*-AATTACTTTTATGTCCCCTGTTGACTGG-3*’*;

### Immunofluorescence, confocal laser microscopy, and Pearson’s coefficient of co-localization

HUVEC were grown on four-chambered slides (Thermo Fisher Scientific). Cells were then washed briefly with PBS and fixed for 20 min rocking in 4% (w/v) paraformaldehyde on ice. Bovine serum albumin (1% in PBS) was utilized to block cells for 1 h, washed thrice in PBS, incubated for 1 h at RT with primary antibodies, washed thrice in PBS, and then incubated for 1 h at RT with secondary antibodies. After washing thrice in PBS, nuclei were stained with DAPI and mounted with a hard set mounting medium (Vector Labs, Burlingame, CA) and coverslips. Using a 63x, 1.3 oil-immersion objective in a Zeiss LSM-78 confocal laser-scanning microscope, fluorescence images were acquired at single optical sections of 1.0 μm, collected with the pinhole set to 1 Airy Unit for all three channels. Line scanning was performed as described previously (Thurston et al., 2012; Goyal et al., 2012). All images were analyzed in ImageJ and Photoshop CS6 (Adobe Systems). Images were acquired in a single plane to determine co-localization. We calculated the weighted Pearson’s coefficient of co-localization using the co-localization function in the LSM-780 Zen software package as described previously (Neill et al., 2016).

### Transient siRNA-mediated knockdown

Transient knockdown of VEGFR2 (Flk-1) in HUVEC was accomplished via transfection of validated siRNA directed against human VEGFR2 (sc-9318; Santa Cruz Biotechnology). Scrambled siRNA (sc-37007; Santa Cruz Biotechnology) was used as control. Gelatinized six-well plates seeded with 2 × 10^5^ HUVEC were grown at 37 °C and 5% CO_2_ until 70% confluent. VEGFR2 or scrambled siRNA duplex (10 pM) were added to Lipofectamine 2000 (Invitrogen) in serum-free OptiMEM (Thermo Fisher Scientific). Cells were washed with PBS, incubated with the OptiMEM solution for 6 h at 37 °C and 5% CO_2_, and incubated overnight with full HUVEC media in the same culture conditions. The media was then replenished, and treatment with vehicle or endorepellin began after 48 h of transient knockdown. Verification of siRNA-mediated knockdown was accomplished via immunoblotting.

### LC-MS/MS analysis of conditioned media

Media of HUVEC treated for 24 h with vehicle, endorepellin (200 nM), or Torin 1 (20 nM) were filtered (0.22 μm), flash frozen in liquid nitrogen, and stored at −80 °C. Unsaturated disaccharide standards of CS (0S_CS-0_: ΔUA-GalNAc; 4S_CS-A_: ΔUA-GalNAc4S; 6S_CS-C_: ΔUA-GalNAc6S; 2S_CS_: ΔUA2S-GalNAc; 2S4S_CS-B_: ΔUA2S-GalNAc4S; 2S6S_CS-D_: ΔUA2S-GalNAc6S; 4S6S_CS-E_: ΔUA-GalNAc4S6S; TriS_CS_: ΔUA2S-GalNAc4S6S), unsaturated disaccharide standards of HS (0S_HS_: ΔUA-GlcNAc; NS_HS_: ΔUA-GlcNS; 6S_HS_: ΔUA-GlcNAc6S; 2S_HS_: ΔUA2S-GlcNAc; 2SNS_HS_: ΔUA2S-GlcNS; NS6S_HS_: ΔUA-GlcNS6S; 2S6S_HS_: ΔUA2S-GlcNAc6S; TriS_HS_: ΔUA2S-GlcNS6S), and unsaturated disaccharide standard of HA (0S_HA_: ΔUA-GlcNAc), where ΔUA is 4-deoxy-α-L-*threo*-hex-4-enopyranosyluronic acid (purchased from Iduron, UK) (Table S1). Chondroitin lyase ABC from *Proteus vulgaris* was expressed in the Linhardt laboratory. Recombinant *Flavobacterial* heparin lyases I, II, and III were expressed using *Escherichia coli* strains provided by Jian Liu (College of Pharmacy, University of North Carolina). AMAC and sodium cyanoborohydride (NaCNBH_3_) were obtained from Sigma (St. Louis, MO, USA). All other chemicals were of HPLC grade. Spectra/Por 6 dialysis tubing (molecular weight cut off: MWCO = 3 kDa) was from spectrum labs (Rancho Dominguez, CA, USA).

Media samples were defrosted at 4 °C and mixed well via vortex. An aliquot of 300 μl of each sample was desalted by 3 kDa spin column and mixed with 300 μl digestion buffer. Recombinant heparin lyase I, II, III and chondroitin lyase ABC (10 mU each) were then added to the reaction buffer and placed in a 37 °C incubator overnight. The reaction was terminated by eliminating enzyme via passing through 3 kDa MWCO spin columns. The filter unit was washed twice with 300 μl distilled water and the filtrate was lyophilized. The dried samples were AMAC-labeled by adding 10 μl of 0.1 M AMAC in DMSO/acetic acid (17/3, v/v) incubating at RT for 10 min, followed by adding 10 μl of 1 M aqueous NaBH_3_CN and incubating for 1 h at 45 °C. The resulting samples were centrifuged at 13,200 rpm for 10 min. Finally, each supernatant was collected and stored in a light resistant container at RT until analyzed via LC-MS/MS. LC was performed on an Agilent 1200 LC system at 45 °C using an Agilent Poroshell 120 ECC18 (2.7 μm, 3.0 × 50 mm) column. Mobile phase A (MPA) was a 50 mM ammonium acetate aqueous solution, and the mobile phase B (MPB) was methanol. The mobile phase passed through the column at a flow rate of 300 μl/min. The gradient was 0-10 min, 5-45% B; 10-10.2 min, 45-100% B; 10.2-14 min, 100% B; 14-22min, 100-5% B.

A triple quadrupole mass spectrometry system equipped with an ESI source (Thermo Fisher Scientific, San Jose, CA) was used a detector. The online MS analysis was at the Multiple Reaction Monitoring (MRM) mode. MS parameters: negative ionization mode with a spray voltage of 3,000 V, a vaporizer temperature of 300 °C, and a capillary temperature of 270 °C. The conditions and collision energies for the all of the disaccharides MRM transitions are listed (Table S2).

### Quantification and statistical analysis

Immunoblots were analyzed via scanning densitometry (ImageJ). Significance was determined by two-tailed unpaired Student’s *t* test. Mean differences were considered statistically significant at P < 0.05.

## Acknowledgements

We thank members of the Iozzo laboratory for helpful discussions. This work was supported in part by NIH grants CA39481 and CA47282 to RVI. Carolyn Chen and Maria Gubbiotti were supported in part by NIH training grants T32 AR052273 and T32 AA07463, respectively.

The authors declare no competing financial interests.

Author contributions: Conceptualization: C.G. Chen, M.A. Gubbiotti, R.V. Iozzo; Methodology: C.G. Chen, M.A. Gubbiotti, R.J. Linhardt, R.V. Iozzo; Validation: C.G. Chen, R.V. Iozzo; Formal analysis: C.G. Chen, R.V. Iozzo; Investigation: C.G. Chen, X. Han, Y. Lu, R.J. Linhardt, R.V. Iozzo; Writing: C.G. Chen, M.A. Gubbiotti, R.V. Iozzo; Visualization: C.G. Chen; Supervision: R.V. Iozzo; Funding acquisition: R.V. Iozzo.

## Online supplemental material

Fig. S1 shows that treatment with endorepellin or Torin 1 has no effect on *HAS2* or *HAS3* mRNA levels and provides additional validation for Fig. 1 and 2. Fig. S2 shows that endorepellin and Torin 1 separately induce co-localization of HAS2 into p62-positive autophagosomes and that autophagic flux evokes HAS2 co-localization with autophagosomal players ATG9A, LC3, and p62. Fig. S3 demonstrates that HAS2 does not complex directly with p62 in co-immunoprecipitation (co-IP) assay and supports the confocal images in Fig. 4. Tables 1 and 2 show the structure of unsaturated disaccharide standards and conditions and collision energies for disaccharide Multiple Reaction Modeling transitions, respectively, and corresponds with the results in Fig. 5.

## Supplementary Figures

**Figure S1.**
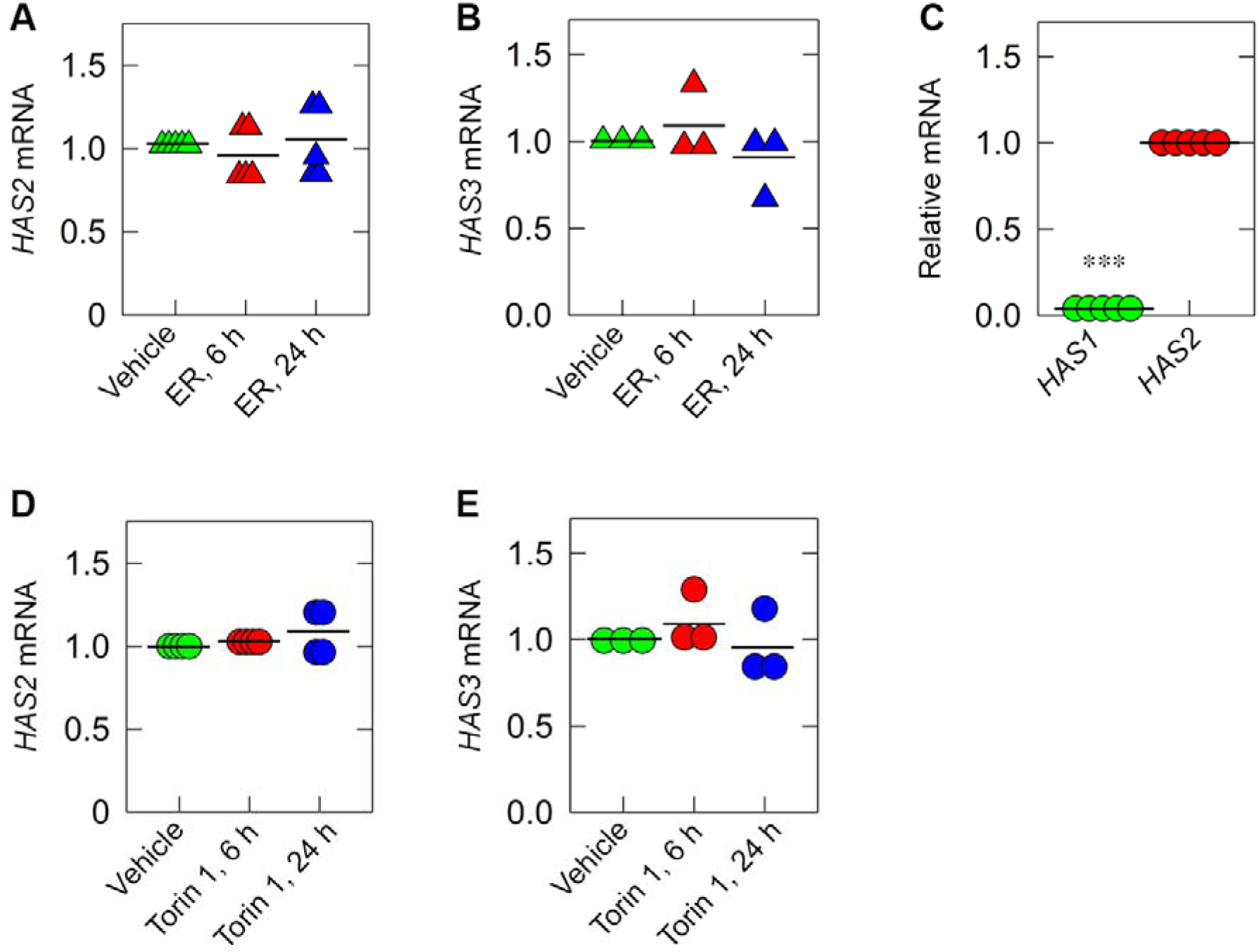
Treatment with endorepellin or Torin 1 has no effect on *HAS2* or *HAS3* mRNA levels. Relative *HAS2* **(A)** and *HAS3* **(B)** mRNA levels in HUVEC treated with endorepellin for 6 and 24 h vis-à-vis vehicle (PBS); *n* = 5 and 3, respectively. **(C)** Relative mRNA levels of *HAS1* and *HAS2* in untreated HUVEC. Relative *HAS2* **(D)** and *HAS3* **(E)** mRNA levels in HUVEC treated with Torin 1 for 6 and 24 h vis-à-vis vehicle (DMSO); *n* = 4 and 3, respectively. Data are presented as the mean (horizontal bar). Statistical analyses were calculated via Student’s *t* test, ***, P < 0.01.

**Figure S2.**
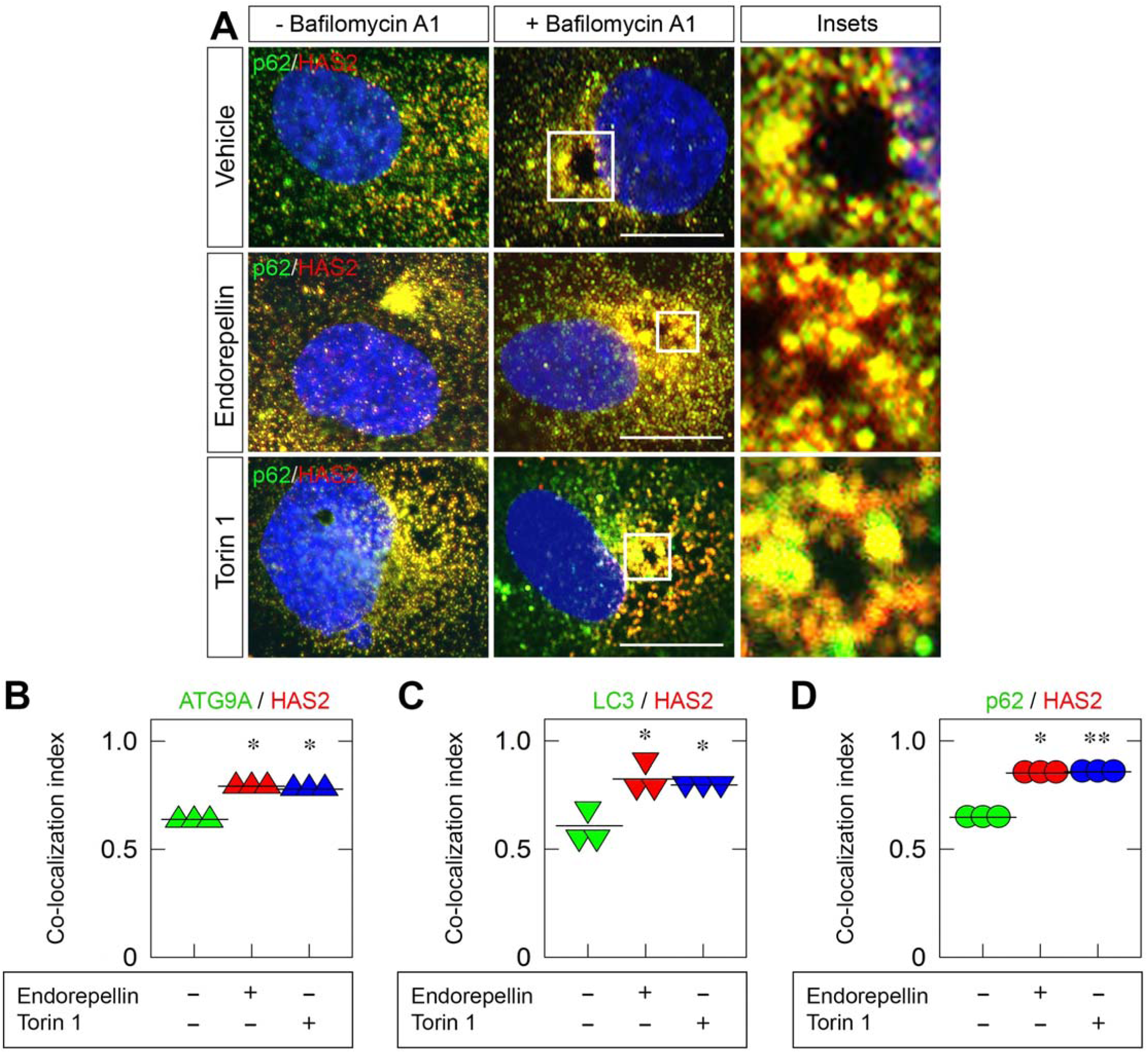
Endorepellin and Torin 1 separately induce co-localization of HAS2 into p62-positive autophagosomes. (A) Immunofluorescent confocal images of HUVEC following treatment with vehicle (DMSO), endorepellin (200 nM) or Torin 1 (20 nM) ±Bafilomycin A1 (500 nM) for 6 h with 1 h pre-treatment of Bafilomycin A1. Cells are dually labeled for HAS2 (red) and p62 (green). Enlarged insets depict co-localized (yellow) signal from cells treated Bafilomycin A1 with vehicle, endorepellin, or Torin 1. Scale bar, 10 μm. Autophagic flux evokes HAS2 co-localization with autophagosomal players ATG9A, LC3, and p62. Pearson’s coefficient of co-localization quantifying signal overlap of HAS2 with **(B)** ATG9A, **(C)** LC3, and **(D)** p62 in immunofluorescent confocal images of HUVEC treated with vehicle (DMSO), Bafilomycin A1 (500 nM, 6 h), endorepellin (200 nM, 6 h), and Torin 1 (20 nM, 6 h). Co-localization of pixels within the entire cell was quantified, and data are presented as the mean (horizontal bar); n=3. Statistical analyses were calculated via Student’s t test. *p<0.05; **p<0.01.

**Figure S3.**
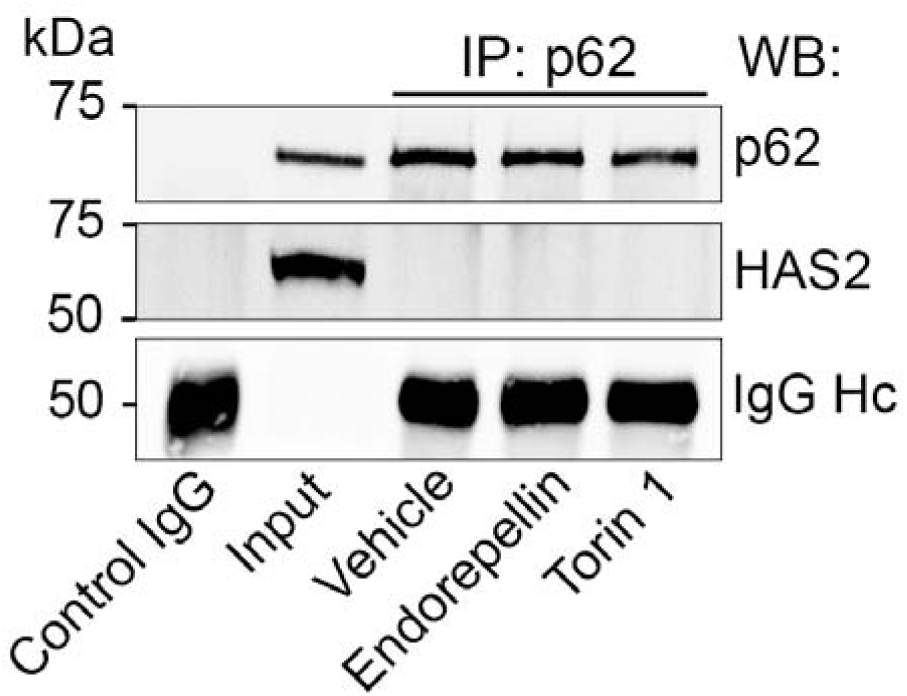
HAS2 does not complex directly with p62 in co-immunoprecipitation (co-IP) assay. Representative co-IP of HUVEC stimulated with endorepellin (200 nM) or Torin 1 (20 nM) for 6 h. The cells were lysed, immunoprecipitated with an anti-p6 antibody and subjected to WB with anti-p62 and anti-HAS2 antibodies as indicated. IgG heavy chain (Hc) is presented as a loading control.

**Table S1.**
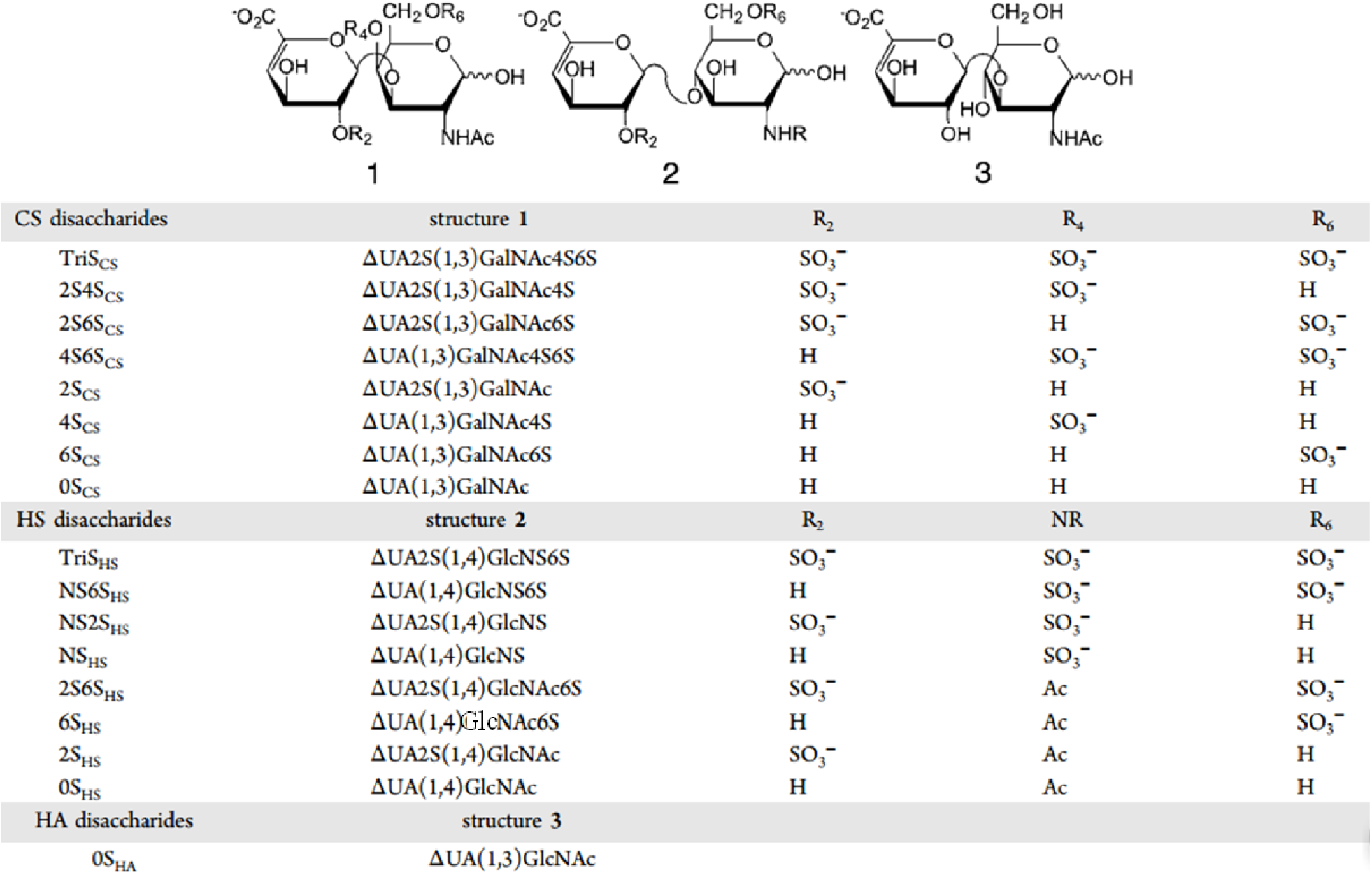
Structure of unsaturated disaccharide standards.

**Table S2.** Conditions a transitions.

| Disaccharide | Precursor ion (m/z) | Production ion (m/z) | Collisional energy (V) | Tube lens (V) |
| --- | --- | --- | --- | --- |
| TriS<sub>HS</sub> | 770 | 690 | 20 | -70 |
| NS6S<sub>HS</sub> | 690 | 610 | 20 | -75 |
| NS2S<sub>HS</sub> | 690 | 610 | 20 | -75 |
| TriS<sub>CS</sub> | 405.8 | 366 | 20 | -70 |
| NS<sub>HS</sub> | 609.8 | 354 | 33 | -100 |
| 2S4S<sub>CS</sub> | 732 | 652 | 20 | -80 |
| 2S6S<sub>HS</sub> | 732 | 652 | 20 | -80 |
| 2S6S<sub>CS</sub> | 732 | 652 | 20 | -80 |
| 4S6S<sub>CS</sub> | 732 | 652 | 20 | -80 |
| 6S<sub>HS</sub> | 652 | 396 | 37 | -110 |
| 2S<sub>HS</sub>/2S<sub>CS</sub> | 652 | 157 | 29 | -106 |
| 4S<sub>CS</sub> | 652 | 396 | 40 | -110 |
| 6S<sub>CS</sub> | 652 | 396 | 31 | -110 |
| 0S<sub>HS</sub> | 572 | 396 | 28 | -90 |
| 0S<sub>HA</sub> | 572 | 396 | 26 | -90 |
| 0S<sub>CS</sub> | 572 | 396 | 28 | -90 |

